# The strength and form of natural selection on transcript abundance in the wild

**DOI:** 10.1101/2020.02.24.948828

**Authors:** Freed Ahmad, Paul V. Debes, Ilkka Nousiainen, Siim Kahar, Lilian Pukk, Riho Gross, Mikhail Ozerov, Anti Vasemägi

## Abstract

Gene transcription variation is known to contribute to disease susceptibility and adaptation, but we currently know very little about how contemporary natural selection shapes transcript abundance. We estimated selection on transcript abundance in cohort of a wild salmonid fish (*Salmo trutta*) affected by a myxozoan parasite through mark-recapture field sampling and the integration of RNA-seq with classical regression-based selection analysis. We show, based on fin transcriptomes of the host, that infection by an extracellular myxozoan parasite (*Tetracapsuloides bryosalmonae*) and subsequent host survival is linked to the upregulation of mitotic cell cycle genes. We also detect a widespread signal of disruptive selection; intermediate transcription levels were frequently associated with reduced host survival. Our results provide insights how selection can be measured at the transcriptome level to dissect the molecular mechanisms of contemporary natural selection driven by climate change and emerging anthropogenic threats.

## Introduction

Understanding how natural selection acts on traits and eventually on organisms represents a fundamental challenge in biology (*1*). Using a now classical regression-based approach (*2*), ecologists have generated thousands of phenotypic selection estimates over the past 35 years; these estimates help to understand the contemporary selection processes in nature and enable comparisons of the strength and mode of selection across traits and species (*3, 4, 5*).

However, despite this wealth of phenotypic selection estimates and a large number of studies that indirectly infer the roles of different evolutionary forces in shaping gene expression patterns (*6, 7*), we know very little about how natural selection affects transcript abundance in the wild (*8*). This is surprising given that variation in transcript abundance is of central importance to evolution (*6–10*), linking molecular functions to performance and Darwinian fitness.

Here, we present an integrative approach investigating how contemporary natural selection shapes transcriptomic variation by combining analyses of selection differentials and gradients (*2*) with the high-throughput screening of molecular phenotypes at the gene transcription level. Such use of the so-called molecular phenotypes has been highly successful in medical science for discovering the mechanisms underlying complex human diseases (e.g., *11, 12*), but we currently know very little about how within-generation natural selection in the wild translates to changes at the RNA and protein levels (*13*). However, regression-based and distributional selection differentials and gradients (*2, 14*), which measure the effect of a trait on relative fitness in standard deviation trait units, can be used to estimate the form and strength of contemporary natural selection on any quantitative trait, including transcript abundances, allowing direct comparisons among traits, populations and species (*2*).

We focus on a host-parasite system consisting of brown trout (*Salmo trutta*) as the host and a myxozoan parasite (*Tetracapsuloides bryosalmonae*), the causative agent of temperature-dependent proliferative kidney disease (PKD) in salmonid fishes (*15*). Recent work has demonstrated that *T. bryosalmonae* is widespread in Europe and North America (*16–20*). At elevated temperatures (>15-18°C), this parasite causes high mortality in wild and farmed salmonids (*21–23*). The parasite has a complex two-stage life cycle in which freshwater bryozoans and salmonid fishes are consecutive hosts (*15*). Mass release of *T. bryosalmonae* spores from bryozoans occurs from spring to early summer (*23*), resulting in the synchronized infection of young-of-the-year fish through skin and gills (*24*). Inside the salmonid host, the parasite induces an inflammatory response and tumor-like chronic lymphoid hyperplasia in the kidney (*23, 25*). The impairment of the kidney, the major organ responsible for hematopoiesis in fish, results in anemia (*23, 26*), which decreases the oxygen transportation capacity, lowering the maximum metabolic rate and upper thermal tolerance (*27*). The reduction of the aerobic and renal capacity, combined with decreased oxygen solubility and increased oxygen demand at higher temperatures, makes PKD a serious threat to cold-water salmonid populations in regions affected by warm summers, which are expected to become more frequent under global warming (*15*). Compared to many other host-parasite systems, brown trout and *T. bryosalmonae* represents a highly suitable model for studying contemporary natural selection on host gene expression in the wild because the parasite shows a high prevalence (*23*) and imposes a strong temperature-dependent effect on host physiology, performance (*27*) and survival (*23*). Furthermore, many challenges associated with field data, such as differences in host age, infection onset, and conspecific coinfection dynamics (*28–29*) or host exposure avoidance (*30*), are minimal or absent.

To quantify the strength and form of within-generation selection on transcript abundance, we collected small fin biopsies from wild juvenile trout in August, when we expected all individuals to be infected, for transcriptome and multilocus fingerprint profiling, after which they were released back into their native environment. Approximately one month later, after the anticipated period of parasite-associated mortality, we recaptured and identified survivors based on multilocus genotype information and tested whether fin-tissue transcript abundances measured in August correlate with survival. To further elucidate the transcriptional signatures linked with the observed mortality, we also measured the *T. bryosalmonae* load in kidney tissue among survivors to identify transcripts and protein–protein interaction (PPI) networks that associate with both survival and parasite load.

## Results

### Parasite load (PL), survival and transcript abundance

Among 18,717 host genes expressed in pelvic fin tissue, 804 covaried with the parasite load (PL) quantified in kidney tissue one month later (unadjusted P<0.01, FDR<0.19; Fig. 1c; Table S2). These results indicate that among the top 804 transcripts, approximately 650 likely represent true positives showing a genuine association between transcript abundance and PL. Consistent with the linear regression results, a discriminant analysis of principal components (DAPC (*31*)) identified a host transcriptome signature predictive of PL (Fig. 1b). Gene Ontology (GO) analysis revealed that the genes positively correlated with PL represent a nonrandom set of genes showing enrichment for 59 GO terms (GOrilla (*32*), FDR<0.05; Table S3), with the top three (FDR<7.7 × 10^−6^) biological processes of cell division (GO:0051301, n = 41), mitotic cell cycle process (GO:1903047, n = 48) and cell cycle process (GO:0022402, n = 57).

**Fig. 1.**
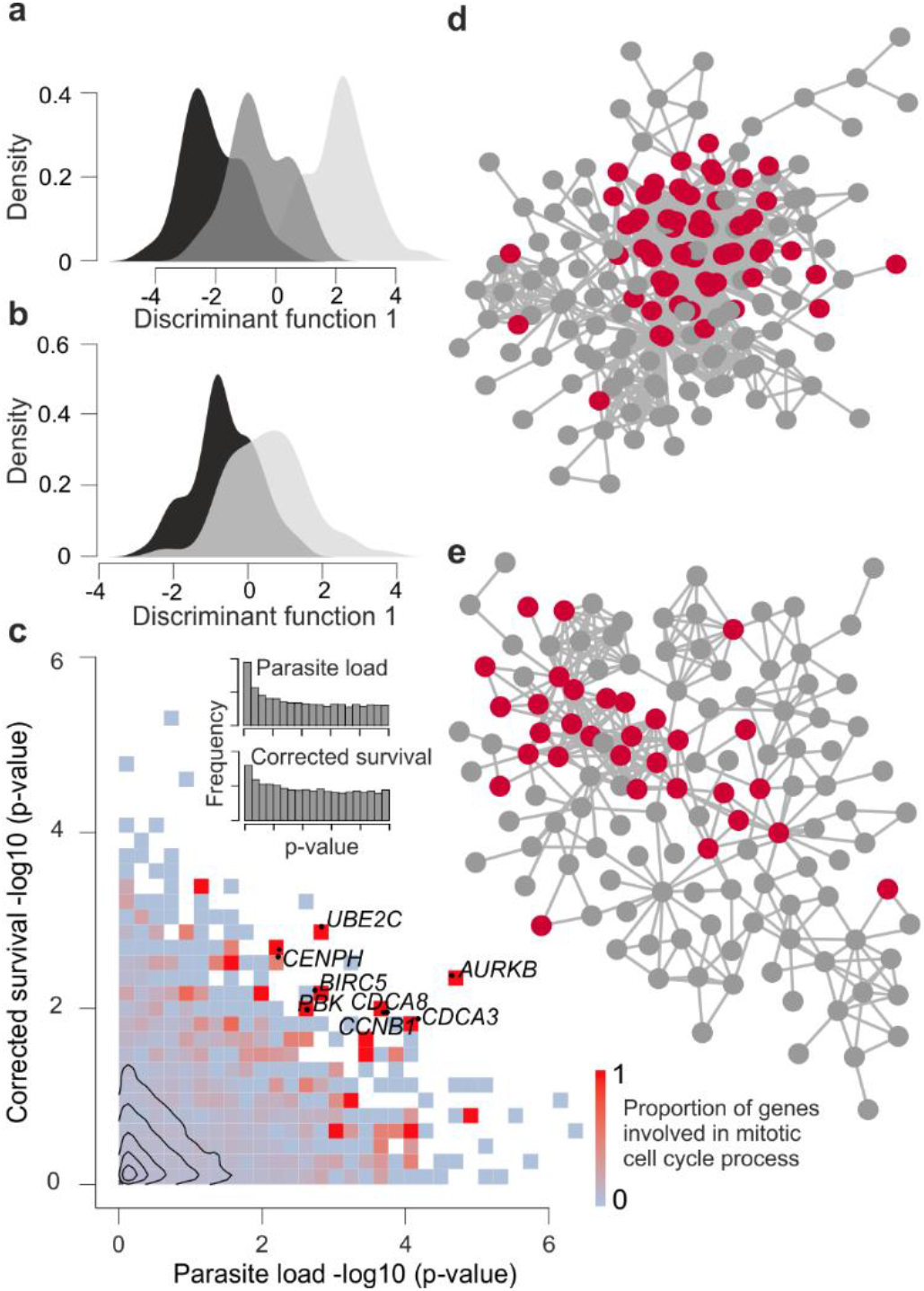
Transcriptome responses in relation to parasite load (PL) and survival. **a** Density distribution of the first discriminant scores corresponding to low, intermediate and high PLs (black, dark gray and light gray areas, respectively). **b** Density distribution of the first discriminant scores corresponding to survivors and nonsurvivors (light gray and black areas, respectively). **c** Proportion of genes involved in mitotic cell cycle presented as a heatmap. The inserted histograms reflect an excess transcripts associated with PL and survival. Contours reflect the density of individual transcripts. Protein–protein interaction (PPI) network with transcripts positively correlated with PL and survival (**d** and **e**, respectively). Mitotic cell cycle genes (GO:0000278) within the PPI networks are shown as red circles.

Fish survival was predicted with 84% accuracy based on the transcription profiles for 1,270 genes using random forest (RF) analysis. Similar to analysis of PL, both DE and DAPC analyses revealed a gene expression signature that covaried with survival (n = 416 genes, unadjusted P<0.01, FDR<0.45; Fig. 1b, 1c; Table S2). These results suggest that among the top 416 transcripts, approximately 229 transcripts likely represented true positives showing genuine associations between transcript abundance and survival.

### Potential links between survival and parasite load (PL)

Protein-protein interaction (PPI) network analysis using String-db indicated that both survival-associated transcripts (PPI enrichment, P<0.001) and transcripts positively correlated with PL (PPI enrichment, P<1.0 × 10^−16^) were highly enriched for genes involved in the mitotic cell cycle (GO:0000278; FDR = 7.45 × 10^−7^ and 3.87 × 10^−24^, respectively; Fig. 1d, 1e). Among the genes associated with both survival and PL are several known oncogenes and tumor suppressors (e.g., *AURKB, BTG1, UBE2C, BIRC5, EEF2K*)(Fig. 1C). On the other hand, survival was not dependent on fish size (Welch Two Sample *t*-test, n = 278, P = 0.216; Wilcoxon rank sum test, P = 0.190).

To further explore the relationships between PL and survival at the transcriptome level, we carried out a weighted gene coexpression network analysis (WGCNA (*33*)). The survival-associated genes clustered into seven gene coexpression networks (Fig. 2a), which included two modules that correlated with PL (Fig. 2b). One particular module consisting of 27 genes (depicted in red in Fig. 2b) showed strong enrichment for the mitotic cell cycle (GOrilla, FDR = 5.9 × 10^−5^; PPI enrichment P<1.0 × 10^−16^), similar to the results from individual transcript analysis. Within the red module, the survival-linked genes that showed the highest correlations with PL were *AURKB, UBE2C, BIRC5* and *CENPH,* which are known key regulators of the mitotic cell cycle (Fig. 2c, 2d).

**Fig. 2.**
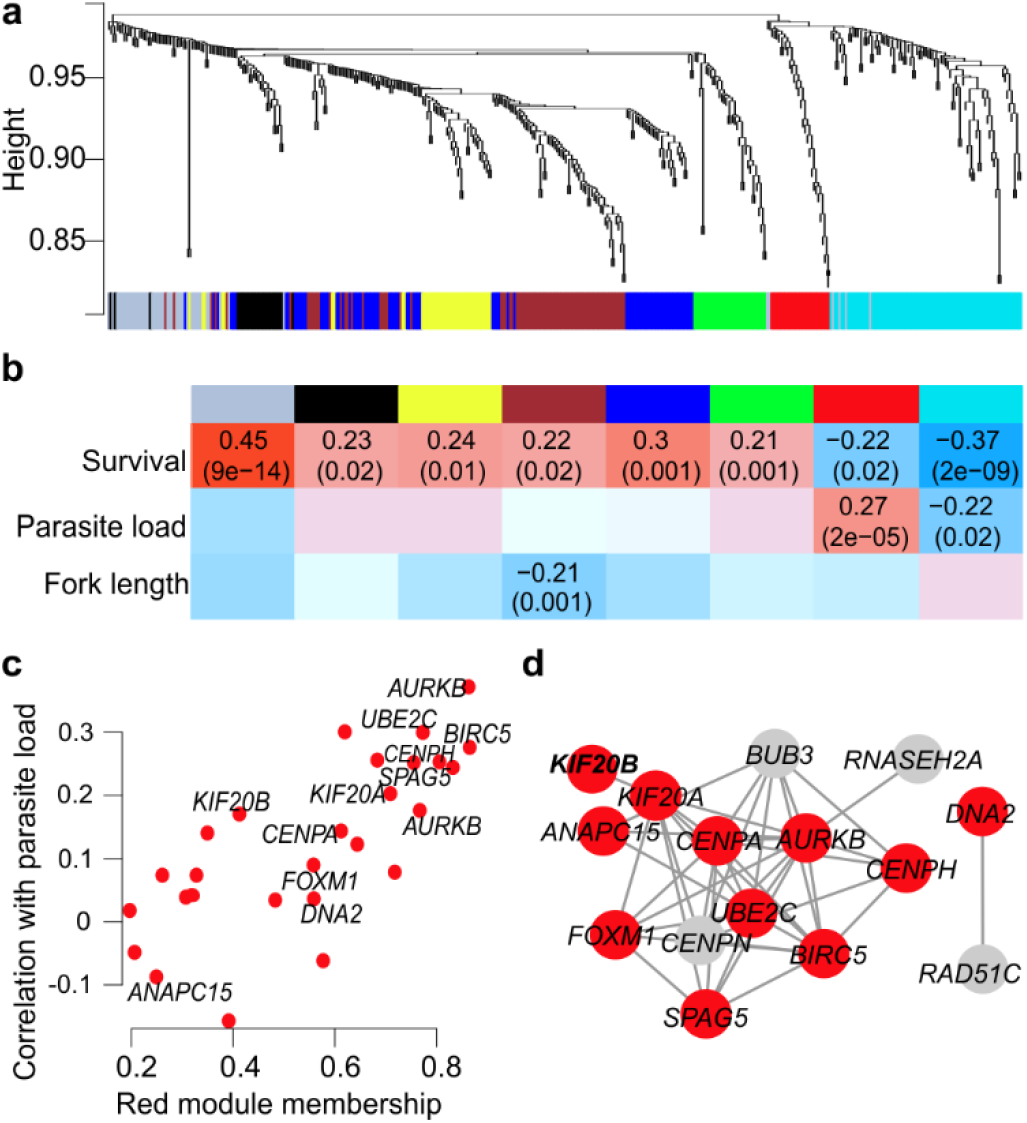
Weighted gene coexpression network analysis (WGCNA) of survival genes and their relationship with parasite load. **a** Gene dendrogram with the corresponding seven modules. Each color represents a module with highly connected genes. **b** Relationships of module eigengenes and survival, parasite load (PL) and fork length (FL). The numbers in the table represent the Pearson correlation coefficients between the corresponding module eigengene and trait, with the p-values in brackets. **c** Module membership of the red module genes and the corresponding Pearson correlation coefficients with parasite load. d Protein-protein network of the red module genes involved in the mitotic cell cycle (GO:0000278) shown as red circles.

### Linear selection differentials and gradients

To quantify the strength and form of contemporary natural selection on transcript abundance, we first estimated standardized linear (*s*) and quadratic selection differentials (*λ*) for 18,717 transcripts. Selection differentials quantify selection (both direct and indirect) on a trait in terms of the effects of trait values on relative fitness in units of standard deviations of the trait, allowing direct comparisons among traits, populations and species (*2*). We compared their magnitudes to a large dataset of phenotypic selection estimates based on a variety of traits and taxa (1,834 published estimates of *s*)(*4*). The vast majority of s values, which measure the change in a population’s mean trait value before and after selection, were small (median(|*s*|) = 0.047; 95% values of *s* between −0.132 and 0.129, *ŝ* ± se = −0.0011 ± 0.0005, 118716 = −2.14, *P* = 0.033), whereas the co-regulated gene modules associated with survival showed larger values of *s* (Fig. 3a). Similar results were obtained for the linear differential estimates calculated using both uncorrected and corrected survival information (Fig. S5).

**Fig. 3.**
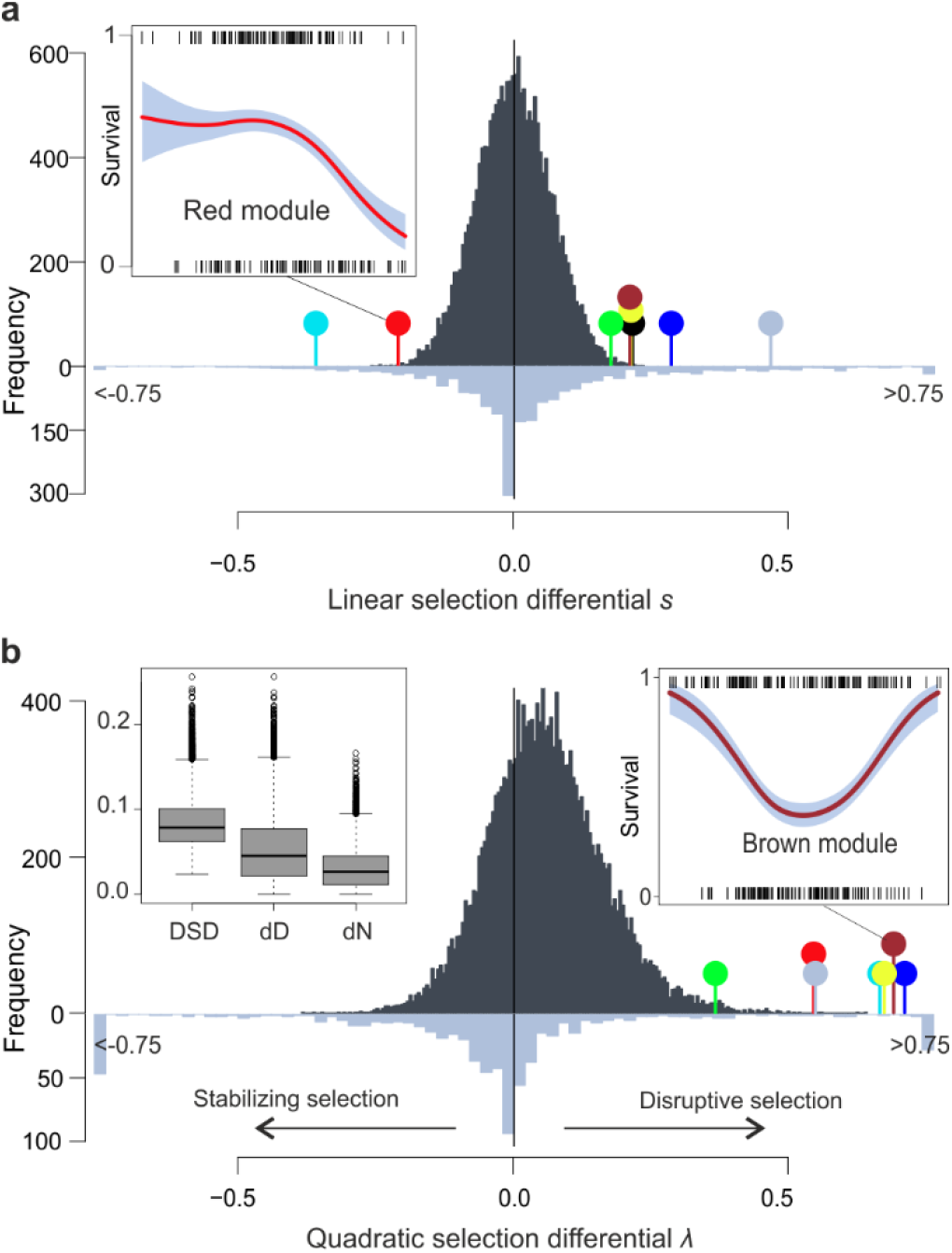
Strength of survival selection on 18,717 transcripts and published phenotypic traits. **a** Linear selection differentials s. **b** Quadratic selection differentials *λ*. Differentials for transcripts and published phenotypic traits (*4*) are shown as dark and light gray histograms, respectively. Negative and positive *λ* values reflect stabilizing and disruptive selection, respectively. Estimates <-0.75 were assigned a value of −0.75, and estimates >0.75 were assigned a value of 0.75. Selection differentials for the WGCNA gene modules are shown as colored pins. The inserted figures illustrate the relationships between survival and module eigengenes as cubic spline (*69*) functions (95% CI in gray) for the red and brown modules; short insert lines reflect individual data. The inserted boxplot illustrates total selection as measured by the distributional selection differential (DSD; *14*), which is broken down into components representing selection on the trait mean (dD = |s|) and selection on the shape of the trait distribution (dN). The line across the box represents the median; the box edges represent the upper and lower quartiles; the whiskers extend to a maximum of 1.5 × IQR beyond the box; and the points represent outliers.

Next, we measured the linear selection gradients (*β*) for each of the 416 transcripts that covaried with survival after removing the effect of indirect selection in a multiple regression framework using principal component scores (*34*). Among the reconstituted linear selection gradients for the 416 DE genes, a total of 67 estimates of *β* remained significant (unadjusted P<0.01, FDR<0.05). Similar to the differentials, genes showing significant linear gradients were enriched for regulation of the cell cycle (GO:0051726, FDR = 3.1× 10^−2^, n = 9) and comprised known key regulators of mitotic cell cycle, including *CENPH, CENPN* and *KIF20A*. Interestingly, body size, which had no direct effect on survival, showed a significant selection gradient (*β* = 0.473; FDR = 6.0× 10^−4^). However, the reconstituted selection gradients for 416 DE genes either controlled or nor controlled for body size (Table S2) were highly correlated (r^2^ = 0.950) indicating that fish size has little effect on these estimates.

### Nonlinear selection

Direct comparison of the strength of linear and nonlinear selection using distributional selection differentials (*14*) revealed that the linear component of selection was generally stronger than the nonlinear component, which represents selection on the shape of the trait distribution (mean dD = 0.053, dN = 0.031; Signed test, P = 9.4 × 10^−206^; Fig. 3b). Nevertheless, for 7,273 (40.8%) genes, the nonlinear differentials were higher than the linear selection differentials (Fig. S6). Furthermore, permutations indicated that while a small proportion of transcripts were affected by directional selection, the dataset was highly enriched for transcripts influenced by disruptive selection, reflecting the elevated survival associated with extreme transcript abundance (Fig. 4); the distribution of *λ* was shifted strongly towards the right tail (Fig. 3b, 95% values of *λ* between −0.137 and 0.279), and its mean differed from zero (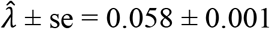, t_18716_ = 78, *P*<2.2 × 10^−16^; compared to 658 phenotypic *λ* estimates (*4*), two-sample Wilcoxon test *P*<2.2 × 10^−16^). Similar results were obtained for the quadratic selection differentials calculated for both corrected and uncorrected survival information (Fig. S5).

**Fig. 4.**
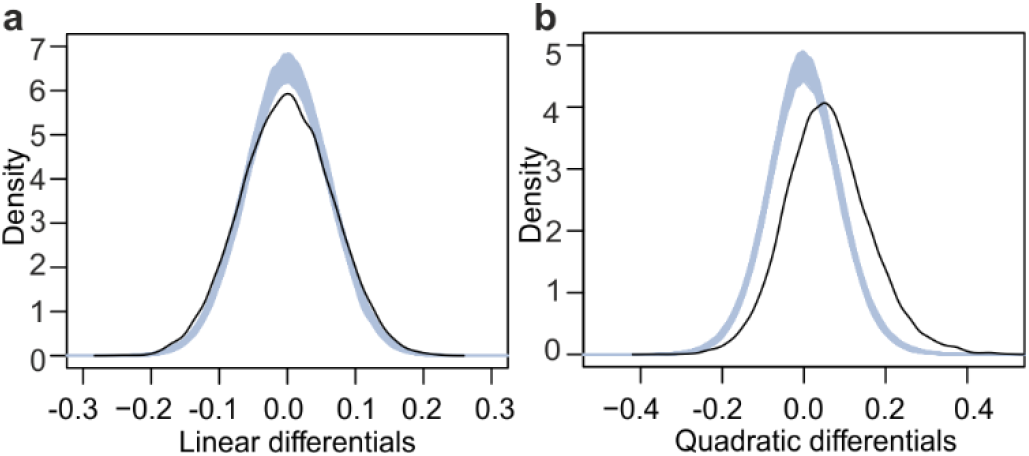
The distribution of linear and quadratic selection differentials. **a** linear selection differentials. **b** quadratic selection differentials. The black line corresponds to observed data, gray lines represent 1000 randomizations (no selection).

Gene Ontology analysis indicated that genes shaped by disruptive selection (λ > 0.2, n = 1,652) showed enrichment of many molecular processes (GOrilla, 51 GO terms, FDR < 0.05; Table S5), including multi-organism process (GO:0051704, FDR = 2.0 × 10^−2^, n = 158), regulation of cell death (GO:0010941, FDR = 4.5 × 10^−2^, n = 187), iron ion homeostasis (GO:0055072, FDR = 3.5 × 10^−2^, n = 18), vesicle-mediated transport (GO:0016192, FDR = 2.8 × 10^−3^, n = 208) and neutrophil activation (G0:0042119, FDR = 4.1 × 10^−2^, n = 73). The transcripts affected by disruptive selection (λ > 0.2) were clustered into six co-expressed gene modules that all showed higher variance among survivors compared to nonsurvivors, a hallmark of disruptive selection favouring extreme trait values (Levene’s test, FDR<2.0 × 10^−4^, Fig. 3b). The constructed co-expressed gene modules showed further enrichment for more specific GO terms, such as the cellular response to cytokine stimulus (brown module, G0:0071345, FDR = 1.5 × 10^−2^, n = 23) and antigen processing and presentation of peptide antigen via MHC class II (brown module, G0:0002495, FDR = 4.97 × 10^−2^, n = 10).

## Discussion

There has long been interest in understanding the relative roles of drift and selection shaping gene expression variation within and between species (*35, 36*). The common approach to this complex question encompasses phylogenetic or comparative analyses that aim to indirectly identify patterns of expression, which do not fit neutral expectations over evolutionarily long time periods. However, these approaches describe the response to selection (R) and not the strength of selection (S) when expressed in the context of the breeder’s equation (R = Sh^2^), where h^2^ is narrow-sense heritability. By combining 3’RNA-sequencing, genetic mark-recapture and selection analysis, we adopted an alternative approach to directly quantify the intensity and form of contemporary natural selection on transcript abundance. As a result, we were able characterize the transcriptomic targets and potential molecular pathways involved in the process of contemporary parasite-driven selection.

Based on uni- and multivariate regression analysis, we identified a small number of transcripts affected by selection. The estimated selection differentials measuring both indirect and direct selection on traits ranged from −0.26 to 0.23, implying that if heritable variation is present and constraints are absent, selection can be strong enough to exert evolutionary changes in transcript abundances at evolutionarily short timescales (*3, 5*). When we compared our transcriptomic data with published phenotypic traits (n = 1,834) in terms of the strength of linear selection (which includes both indirect and direct selection components), similar frequency distributions were observed, with the large majority of estimated differentials being close to zero. However, the phenotypic traits possessed longer “tails” for selection differentials than the transcripts, suggesting either rare but very strong linear selection on some phenotypic traits and/or potential bias due to small sample sizes. On the other hand, selection differentials quantify selection considering each trait separately and measure the total selection acting on the trait (both direct and indirect). Thus, when traits are highly correlated, as is likely among the many transcripts, of which many may be co-regulated, it becomes impractical to distinguish separate influences of individual transcript abundances on relative fitness. To overcome this limitation, we subsequently calculated selection gradients (*β*) for 416 transcripts that covaried with survival to quantify the strength of direct selection on individual genes after removing indirect selection from other correlated transcripts. Altogether, we identified 67 significant *β* estimates ranging from −0.47 to −0.15 and from 0.15 to 0.63, indicating that direct selection on transcript abundances has the potential to cause substantial evolutionary changes at relatively short timescales.

Direct comparison of the strength of linear and nonlinear selection using the distributional selection differentials (*14*) revealed that for 40% of transcripts, the nonlinear differentials were higher than the linear selection differentials. Furthermore, when nonlinear selection was partitioned into stabilizing and disruptive components, our dataset was highly enriched for transcripts showing signatures indicative of disruptive selection. This is unexpected because disruptive selection is thought to be rare in nature (*3, 5*). Moreover, this finding contrasts with the expectation that stabilizing selection is more common than disruptive selection if most populations are well-adapted to their current environment (*3*). On the other hand, it has been suggested that disruptive selection may be more widespread than previously thought, reflecting density-dependent or frequency-dependent competition for resources (*5*). Thus, our results corroborate with phenotypic selection estimates, and also suggest that host transcript abundance may be influenced by disruptive selection in response to parasite infection. For example, it may be more beneficial for a host to either invoke a strong immune response (i.e., highly resistant hosts with the lowest PL and lowest disease expression as measured by kidney hyperplasia) or tolerate the damage from a high PL than to partially control the parasite load (i.e., hosts suffering from damage by both parasites and immunopathology). The functional categorization of genes and gene modules under disruptive selection supported this hypothesis, because they were highly enriched for biological processes related to host immune defence, host-pathogen interactions, cellular repair and maintenance. These inferred functions presumably reflect the complex nature of host-parasite interactions, as the transcripts shaped by disruptive selection were involved in a wide range of molecular processes.

Variation in transcript abundance, similar to that in morphological or physiological traits, is expected to be shaped by selection through whole-animal performance, which can be defined as how well an organism accomplishes certain ecologically relevant tasks (*37*). Therefore, it is pertinent to ask what performance traits are “visible” to selection in the studied host-parasite system. The functional categorization of genes and correlated gene modules provides some clues to this question, as both survival- and PL-associated transcripts shaped by linear selection were highly enriched for genes involved in the mitotic cell cycle. First, we rejected the idea that the transcript variation associated with survival reflect variation for general stress response of the host caused by *T. bryosalmonae* infection. This is because most stress factors lead to an arrest of mitosis (*38–40*), whereas we detected that parasite load (PL) associated with upregulation, rather than downregulation (what may be expected for arrest of mitosis), of the key mitotic cell cycle host genes (*AURKB, UBE2C, BIRC5;* Fig 2). Second, the observed associations between fin tissue transcriptome, PL and survival may reflect the host response to parasite entry because *T. bryosalmonae* enters its salmonid host through surface tissues, which may include fins (*24*). However, we currently lack experimental evidence that *T. bryosalmonae* entry causes upregulation of cell-cycle activity in fin or/and other mucosal tissues of the host. Third, the coupling of the transcription of mitotic cell cycle genes in fin, PL and survival may reflect the severe physiological impact of PKD on the host at a larger organismal level. For example, earlier studies in salmonids have demonstrated that one of the main PKD symptoms is tumour-like proliferation of the lymphoid renal tissue, where the kidney may increase in size to over ten times its normal volume (Fig. S2; *23, 25, 26*). Similarly, PKD causes enlargement of the spleen, and several studies suggest that PKD in salmonids is a systemic disease that affects multiple organs and tissues (*15, 23–27*). In teleosts, pelvic fins consist of epidermis, bony rays, ligaments, nerve fibres, connective tissue cells and blood vessels. The observed associations between cell-cycle related host genes, PL and survival may, therefore, reflect the importance of blood homeostasis and sustaining normal kidney function. However, analysis of multiple tissues, including renal, blood and fin transcriptomes, during the progression of PKD is clearly needed to further describe the molecular mechanisms of the host response, as we currently lack comprehensive understanding of the inflammatory, mitotic and immune processes across organs (*41*). Regardless of the specific physiological mechanism, this work adds to the increasing body of work showing that parasites can influence the host’s cellular machinery (*39, 40, 42*).

Two limitations in this study may be addressed in future research. First, despite the high electrofishing effort, low dispersal and relatively high recapture probabilities, we probably did not recover all survivors. We therefore carried out functional and selection analysis based on both initial recapture information and by treating 13 putatively uncaptured individuals as survivors, as suggested by their transcript profiles that matched recaptured individuals.

However, given that the main findings (e.g. distribution of the linear and nonlinear selection coefficients, gene ontology enrichment patterns) remained very similar irrespective of the classification, imperfect classification of small number of survivors most likely had small effect on the main conclusions. Second, even though it was not possible to analyse the primary target tissue of the parasite (kidney), requiring lethal sampling and thereby preventing survival estimates, the systemic nature of PKD conceivably enabled us to acquire biologically meaningful information from fin biopsies with a minimal expected effect on fish survival (*43*). Similarly, transcript abundances in fin tissue have recently been associated with aging-related mortality in fish, demonstrating the utility of fin tissue for linking gene expression and whole-organism performance (*44*). More generally, because of their major role in pathogen defence, mucosal surface tissues have been widely used to study innate and adaptive immune responses in teleost fishes (*45*).

In summary, our work demonstrates the power and challenges of integrating non-lethal sampling and transcriptomics with classical ecological methods to dissect complex high-order organismal traits, such as survival in the wild, into functionally interpretable molecular processes. As such, our study provides a novel perspective for studying contemporary selection at the suborganismal level and is readily applicable to other species and systems, where non-lethal sampling of blood, mucosal and other tissues is feasible. We anticipate that the approach described here will enable critical information on the molecular mechanisms and targets of natural selection to be obtained in real time, as wild populations increasingly contend with novel selective pressures (*46*), including those imposed by global warming (*47*).

## Methods

### Study population, temporal abundance and temperature records

The coastal river Altja (length 17.6 km, catchment area 46.1 km^2^) flows into the Gulf of Finland in the Baltic Sea (Fig. S1) and supports a small wild anadromous brown trout population with a high prevalence of *T. bryosalmonae* (*18, 26, 17*). Records on young-of-the-year trout abundance in the river Altja across 13 years (2005-2017) was obtained from the Estonian Ministry of Environment Fisheries Monitoring Program. River water temperature was measured twice per day (at noon and midnight) over two years (2014 and 2015) using automatic temperature loggers (HOBO 8K Pedant Temperature/Alarm Data Logger, Onset Computer Corporation). Mean monthly air temperature records over a 73-year period (1945-2017, Kunda Coastal Meteorological station, N 59°31′7″, E 26°32′29″, 25 km from river Altja) were obtained from the Estonian Weather Service (Environmental Agency). The studied population showed a strong negative correlation between the mean summer air temperature and young-of-the-year density as well as pervasive temperature dependence of disease severity, consistent with experimental work (*25*) (Fig. S2).

### Field sampling, phenotyping and genetic mark-recapture analysis

On 30 August 2015, we electrofished 278 young-of-the-year trout in the river Altja from the same five areas along a 330 m stretch that were sampled in 2014 (Table S1, Fig. S1, area IDs 1-5). Individuals were anesthetized with buffered MS-222 (SigmaAldrich, St. Louis, MO) and measured for fork length (±1 mm) as a measure of body size. After small biopsies of the right pelvic fin tissue for genetic mark-recapture analysis and 3’ RNA sequencing (see below), we released the recovered trout within their original capture area. As fins regenerate in teleosts, a small fin biopsy is unlikely to impair fish survival (*43*). Biopsies were immediately frozen in liquid nitrogen and stored at −80°C. We used a fin subsample for DNA analysis and individual identification (see below). Approximately one month after initial electrofishing (22-27 Sept), we caught 685 young-of-the-year trout along a 780 m stretch that included the initial 330 m stretch (Table S1). The five initial catch areas (area IDs 1-5) were electrofished thrice (Table S1), and we estimated the capture probability and total number of fish using the depletion method (*48*) implemented in the FSA (Fisheries Stock Assessment) package v. 0.8.17 (*49*) in R v. 3.3.3 (*50*). A high recapture probability in all areas (average catchability 0.65, 95% CI 0.60-0.70) combined with low inferred fish dispersal (based on electrofishing of the extended areas up- and downstream; Extended data Table 1) indicated that only few survivors were not recaptured in September (n = 13.9, 95% CI 7.2-23.1). Among the 685 fish caught in September, we euthanized 363 via MS-222 overdose. The sampling procedure, microsatellite genotyping, measurement of phenotypic traits (fork length, FL as a measure of body size; hematocrit, Hct; kidney-to-muscle ratio as a measure of kidney swollenness, KS) and quantification of the parasite load were carried out as previously described (*16*). The relationships between PL and PKD-linked phenotypic traits (Hct, KS, FL) were analyzed using general linear mixed models in ASReml-R 3.0 (*51*). To control for genetic variation in the expression of the traits, the models accounted for the genetic relationship matrix of the individuals (A) via linking (A^−1^) to random individual effects. The relationship matrix was obtained by genotyping the sampled individuals (see next section) and then reconstructing their parentage using Colony v. 2.0.6.5 (*52*).

### Microsatellite analysis and identification of individual recaptures

We genotyped individuals using 14 highly variable microsatellite loci as previously described (*16*). Individuals with at least 10 successfully genotyped loci were included in the analysis. Recaptured individuals were defined as having identical genotypes with at most one allele mismatch (at least a 95% match when 10 loci were genotyped) using Microsatellite Toolkit (Stephen D.E. Park, Dept. of Genetics, Trinity College). To estimate genotyping error rates, we amplified and genotyped 440 randomly selected samples twice, which indicated low error rates (mean allelic dropout rate: 0.0107, range 0.0013-0.0292; mean false alleles rate: 0.0027, range 0.0010-0.0176).

### Quantitative Real-Time PCR (qPCR)

Parasite load (PL) was determined from kidney tissues collected in September 2015 by quantitative PCR (qPCR) using previously described protocols (*16*). For each sample, we quantified two DNA sequences per run: a 166 bp fragment of *T. bryosalmonae* 18S rDNA sequence (GenBank Accession U70623) and 74 bp fragment of the *S. salar* nuclear DNA sequence (GenBank Accession BT049744.1). Simultaneous quantification of both DNA fragments enabled us to standardize the amount of parasite DNA relative to brown trout DNA. We ran 10 plates (384-well format) on QuantStudioTM 12K Flex Real-Time PCR System (Thermo Fisher Scientific). Each 10 μL reaction contained 200 nM of each primer, 1x HOT FIREPol^®^ EvaGreen^®^ qPCR Supermix master mix (Solis BioDyne, Tartu, Estonia) and 2.5 μL of template DNA (10 ng/μL). Each sample was run in quadruplicate per gene and included four non-template controls per gene to detect possible contamination. To determine the quantification cycle (Cq), we used the online tool Real-time PCR Miner (*53*). Parasite load (PL) was defined as the difference between the Cq values (on the log2 scale) of the two target genes (Cq_*S. trutta*_ Cq_*T. bryosalmonae*_ lower values indicate low PL). Our 2015 qPCR results were calibrated to 2014 results using ten 2014 samples that we repeated along with the 2015 samples (linear regression, PL2014 = −0.193 + 1.031 × PL2015). To estimate technical bias among plates, we included a log10 dilution series (50, 5, 0.5, 0.1 and 0.05 ng/μL) from a reference sample in quadruplicate per gene on every plate. Standardized amplification efficiency was calculated among plates, using plate- and gene-specific efficiencies estimated from the Cq vs. log10-dilution slopes (*16*). Subsequently, we fitted a linear mixed model to estimate parasite load (PL) for each individual that accounted for technical bias among plates and wells (*16*).

### RNA isolation and library preparation for Illumina sequencing

Total RNA was successfully extracted from pelvic fin tissue of 238 individuals (85.6%) out of 278 collected in August 2015 (i.e. survivors and non-survivors) using NucleoSpin^®^ RNA kit (Macherey-Nagel, Düren, Germany) and ensuring RNA quality using the Agilent 2100 Bioanalyzer. We prepared a barcoded library using Lexogen QuantSeq 3’ mRNA-Seq Library Prep Kit FWD for Illumina according to manufacturer’s recommendations (Lexogen, Vienna, Austria). QuantSeq provides an efficient and cost-effective protocol for generation of strand-specific next-generation sequencing libraries close to the 3’ end of polyadenylated RNAs (*54*). This approach is analogous to other tag-based RNAseq approaches, such as TagSeq (*55*), which have been shown to generate more accurate estimates of transcript abundances than standard RNAseq with a fraction of the sequencing effort (*56*). We inspected all barcoded libraries for quality with Fragment Analyzer (Advanced Analytical, AATI) using High Sensitivity NGS Fragment Analysis Kit. The three pooled barcoded libraries, consisting of 64, 91 and 96 individuals, were single-end sequenced using Illumina HiSeq2500 in 14 lanes. For the first two pooled libraries, we generated 125 bp long reads in two lanes. For the remaining twelve lanes, we generated 100 bp reads. Overall, we obtained 2.21 billion raw reads, with 1.5 to 34.6 million reads per individual (median 8.9 million reads). In addition, conventional Illumina mRNA paired-end sequencing (100bp) was carried out for two fin-clip mRNA pools both consisting of seven individuals from the River Altja (29.5 and 25.9 million reads). The data have been deposited with links to BioProject accession number PRJNA517427 in the NCBI BioProject database (https://www.ncbi.nlm.nih.gov/bioproject/). The library preparation for this purpose was done according to Illumina TruSeq^®^ Stranded mRNA Sample Preparation Guide. For adapter trimming and read preprocessing, we used Trimmomatic (v. 0.35 (*57*)) (parameters: ILLUMINACLIP = TruSeq3-SE.fa; HEADCROP = 10; SLIDINGWINDOW = 4:20; LEADING = 5; TRAILING = 5; MINLEN = 40). A total of 44.6 million quality-controlled paired-end reads were retained (23.8 and 20.8 million reads per pool). For the QuantSeq, we used Trimmomatic with slightly different settings (HEADCROP = 12 and MINLEN = 70). We subsequently used Cutadapt (*58*) (version 1.10) to inspect and trim longer runs of poly-As at the end of the QuantSeq reads (parameters: q = 10; b = A{20}; b = A{30}; m = 40) and discarded sequences shorter than 40 bp. We assessed the quality of reads before and after trimming using fastqc (v. 0.11.2). A total of 1.1 to 27.1 million QuantSeq reads per sample remained after quality filtering (median 6.8 million reads).

### Trout fin-specific splice sites, reference genome modifications and mapping

To identify brown trout fin-specific splice sites, we mapped quality-filtered paired-end reads from two pooled fin libraries and three 3’ mRNA-Seq samples with the highest read depth (total 58.4 million reads) to the salmon genome using Hisat2 (v.2.1.0 (*59*)). Full-length transcripts expressed in the trout fins, were subsequently assembled using Stringtie (*60*), and the splice-site locations of these transcripts were extracted using the extract_splice_sites.py script provided with Hisat2. The Atlantic salmon reference genome (GCF_000233375.1) was modified for alternative alleles identified from the paired-end trout RNA reads. We used splice sites as a guide during Hisat2 reference-index building of the modified reference genome, and all quality-controlled reads were aligned from QuantSeq 3’ mRNA-Seq to this reference genome using Hisat2.

### Estimation of transcript abundance and batch effect correction

All alignment data were loaded into R as the RangedSummarizedExperiment object returned by summarizeOverlaps. The original salmon-genome-annotation GFF file was used to dissect exons on the basis of gene information, and union mode was selected for assigning the reads within the exons while considering strand information. Read counts from separate lanes, runs and replicate files were summed to individual counts using collapseReplicates implemented in the R package DESeq2 (*61*). The resulting data object contained a raw read count matrix and phenotypes for each sample. For subsequent analysis, we selected only protein-coding, nuclear genes with an average of > 10 reads per individual. The raw read counts were converted into quantile-normalized log2-counts per million (logCPM) using the voom method (*62*) implemented in the limma package (*63*). Last, pooled library batch effects were removed by employing the ComBat function implemented in the SVA package (*64*), and corrected gene counts were used in differential gene expression analysis.

### Differential expression (DE)

Corrected gene counts were used for testing for the associations of genes with the continuous phenotypic traits (FL in August and PL, KS, and Hct in September) by fitting generalized linear models with each gene as a separate predictor. We calculated Q-values using the qvalue function as implemented in R. To identify genes associated with survival, we iteratively ran a random forest (RF) classification implemented in the R package ranger (*65*) with 100,000 trees in the gene expression matrix and excluded genes with < 0 permutation importance for the next iteration. After recursive runs, the remaining set of 1,270 genes classified individuals into survivors and nonsurvivors with a 16% error rate. Mark-recapture analysis of the three consecutive electrofishing rounds similarly indicated that only a small number (n = 13.9, 95% CI 7.2-23.1) of surviving fish that were sampled in August were not recaptured in September (Table S1). Furthermore, differential gene expression (DE) analysis based on uncorrected survival produced a hill-shaped, rather than uniform, p-value distribution, indicating that misclassification of individuals likely resulted in violation of the statistical test assumptions (Fig. S3). For subsequent analysis, we therefore treated the 13 putatively uncaptured individuals as survivors based on their transcript profiles (Table S4), but the conclusions remained unchanged irrespective of the classification (Fig. S3-S5). For example, linear and non-linear selection estimates were very similar using both uncorrected or corrected survival information (Fig. S6). Also, we observed considerable overlap (n = 171) among the top 416 genes (Fig. S3) between the top lists of observed survivors and corrected survivors, which both showed highly significant enrichment for mitotic cell cycle genes (GO:0000278, Fig. 1, S3). Thus, corrected survival status was used in the differential gene expression analysis using DESeq2 with raw read counts along with the three pooled library IDs as a covariate.

### Discriminant analysis of principal components (DAPC)

We performed DAPC on the gene expression matrix using the dapc function implemented in the R package adegenet (*66*). As the DAPC function requires categorical data, PL values were split into three groups: low (PL < −1.27, n = 24), intermediate (PL = −1.27 – 0.58, n = 39) and high (PL > 0.58, n = 48) using the cut command in R.

### Weighted gene coexpression network analysis (WGCNA)

Genes associated with survival were subjected to automatic network construction and module detection using the blockwiseModules function implemented in the R package WGCNA (*33*). The cutoff for the minimum scale-free topology-fitting index was set to 0.8 (power = 7), and we used biweight midcorrelation (bicor) to estimate correlations (other parameters: networkType = “signed”, minKMEtoStay = 0.2). For the analysis of quadratic selection differentials (see below), similar parameters were used; the lower cutoff for the minimum scale-free topology fitting index was, however, changed to 0.5 (power = 3).

### Gene Ontology (GO) and protein–protein interaction network analysis

Atlantic salmon ortholog gene symbols, entrez IDs, and descriptions in humans (86.8%), zebrafish (3.6%) or other organisms (9.6%) were searched using complete gene names in NCBI using a custom R script (available upon request). GO-enrichment analysis was performed using String-db (*67*) and GOrilla (*32*), in which all ortholog gene symbols were used as a background list. For String-db protein-protein interaction analysis, we used single lists of gene symbols against human protein references (minimum interaction score: 0.70; text mining disabled).

### Quantification of linear and nonlinear selection differentials

We estimated linear and nonlinear (i.e., quadratic) selection differentials for each of the 18,717 gene transcripts quantified in 238 individuals based on both corrected and uncorrected survival (binary status, nonsurvivor = 0, survivor = 1; Table S2). Subsequently, estimates of linear or quadratic selection differentials were computed using generalized linear models under the glm function in R. These models used logit-link functions and binomial error distributions for the binary survival response and the predictor of the mean-centered and variance-scaled (mean = 0, SD = 1) transcript levels (linear differentials) or added a quadratic scaled transcript-level predictor (nonlinear differentials). To transform the logistic regression model coefficients to selection differentials on the relative fitness scale that is meaningful to microevolutionary studies, we used the method suggested by Janzen and Stern (*68*). P-values for each selection differentials were estimated using *t*-tests that were constructed based on logistic regression coefficients, their standard errors, and model residual degrees of freedom. In addition, to calculate the distribution of linear and nonlinear selection differentials when no selection is acting on transcripts, we randomized the survival values (1000 permutations) and estimated selection differentials as described above. To compare the strength of directional and nondirectional selection, we used a recently developed unified measure, the distributional selection differential (DSD (*14*)) for standardized trait values (mean = 0, SD = 1). This enabled us to use a single framework to partition total selection (DSD) into selection on the trait mean (dD) and selection on the shape of the trait distribution (dN).

### Quantification of linear selection gradients

The linear selection gradients for the differentially expressed genes were reconstructed from a subset of principal components (*34*), as this approach not only enables the multicollinearity between the predictors to be handled but is also suitable for cases in which the number of predictors exceeds the number of observations. For this purpose, the principle components were calculated from the correlation matrix of the standardized values of 416 differentially expressed genes, and the linear selection gradients were subsequently computed for the first 55 PCs (explaining 76% of the variation) with the glm function in R, using the logit-link function and binomial error distribution. The eigenvectors from the original 55 PCs were then matrix multiplied with the resulting linear selection gradients to reconstruct the selection gradients for individual genes (*35*). Similarly, the selection gradients for 416 DE genes were calculated by including fork length (FL) as a predictor. The standard errors were reconstituted for these gradients as described in (*35*). The t.statistic was calculated by dividing the gradients with the standard errors, and the P-values were estimated from the results using 237 degrees of freedom. The P-values were corrected for FDR using the p.adjust function in R.

## Supporting information

GOrilla and StringDB Gene Ontology enrichment results

## Data availability

Data supporting the findings of this work are available as Supplementary Information. The sequence data have been deposited with links to BioProject accession number PRJNA517427 in the NCBI BioProject database.

## Code availability

The R code is available upon request.

## Acknowledgments

We thank M.F. Oleksiak, J.M. Henshaw, T. Aykanat, V. Kisand, C. Primmer, T. Tenson, R.J. Pawluk and A. Krasnov for commenting on earlier drafts of the manuscript; K. Haugjärv for help during sample collection; K. Salminen from the Center of Evolutionary Applications, University of Turku for RNA extractions and library preparations; Põlula Fish Rearing Centre (RMK) for their support during fieldwork. Bioinformatic analyses used resources at the CSC—IT Center for Science, Finland. This work was supported by the Academy of Finland (grant 266321), the Estonian Ministry of Education and Research (institutional research funding project IUT8-2), the Estonian Research Council grant (PRG852), Ella & Georg Ehrnrooth foundation, Archimedes Foundation Scholarship in smart specialization growth areas, and a German Research Foundation Research Fellowship (DE 2405/1-1).

## Author contributions

A.V. conceived the study. A.V., F.A., P.V.D., S.K. and L.P. collected the samples. P.V.D. measured hematocrit and kidney swollenness. I.N. carried out microsatellite analysis and parasite quantification. M.O. carried out mark-recapture analysis. F.A. carried out bioinformatic and transcriptomic analyses. F.A. and P.V.D. estimated selection differentials. All authors participated in interpretation of the results. A.V., F.A. and P.V.D. drafted the manuscript, and all others commented. Competing interests: The authors declare no competing interests.

## Materials & Correspondence

Requests and correspondence should be addressed to A.V.

