## Supplementary Materials for "The strength and form of natural selection on transcript abundance in the wild"

**This PDF file includes:**

Figs. S1 to S6

Tables S1 and S4

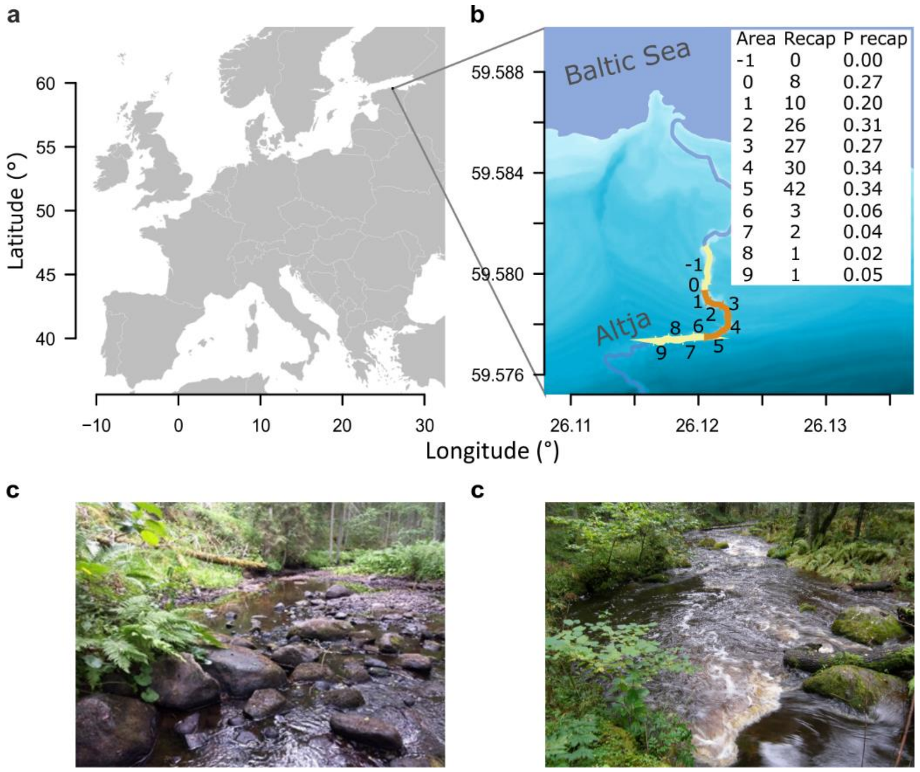

**Fig. S1. Studied field site.** (**a**) Location of the Altja river. (**b**) Areas that were electrofished once and three consecutive times in September are marked with yellow and orange, respectively. Only orange area was electrofished in August. The table insert indicates the number and recapture frequency of fin-clipped fish for each area in September; Photographs from area 0 (**c**) and area 4 (**d**) taken during low- and high-water periods, respectively.

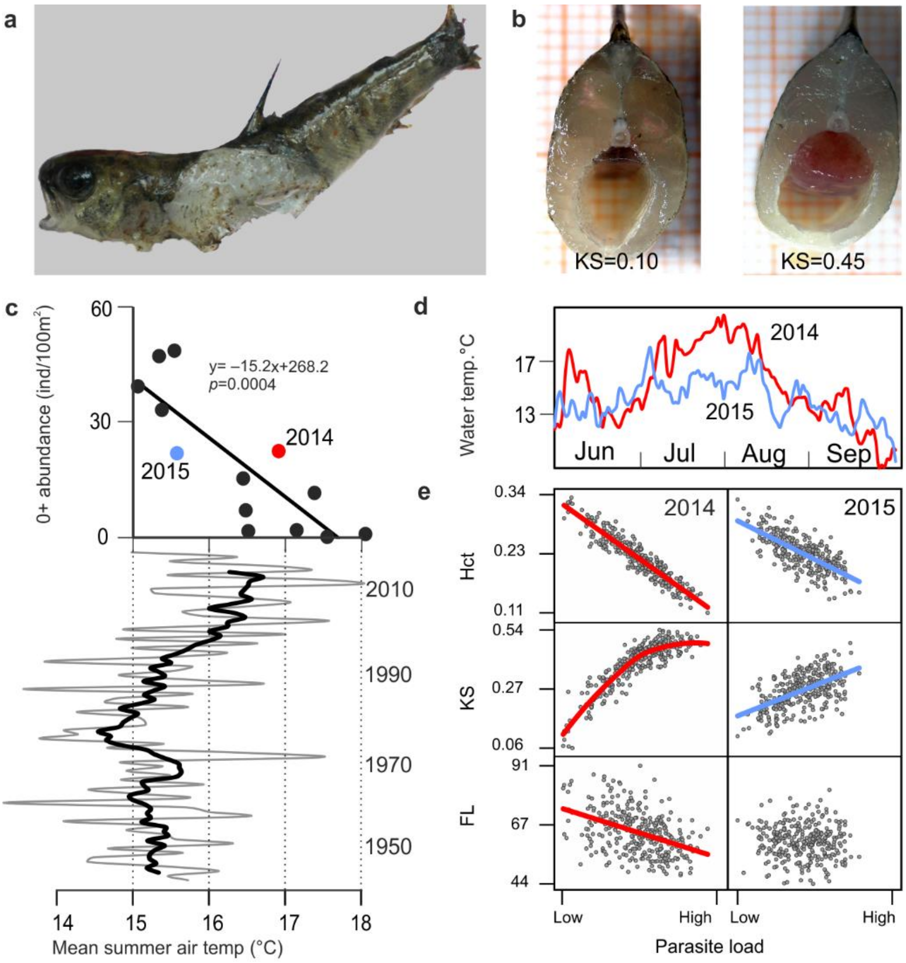


**Fig. S2. Temperature-dependence of PKD in wild trout.** (**a**) Dead young-of-the-year brown trout found in the Altja river with putative PKD-associated death symptoms (swollen kidney, a wide open mouth and flared gills suggestive of anemia); (**b**) Body section of trout with normal (left) and swollen (right) kidney; (**c**) Effect of temperature on juvenile trout abundance during 2005-2017 in the Altja river in relation to average summer air temperature (7-year moving average mean summer air temperature over 73 years is highlighted in bold); (**d**) Water temperature variation over a 4-month period in 2014 (red) and 2015 (blue) in the Altja river; (**e**) Relationships between parasite load (PL) and fork length (FL), kidney swollenness (KS), and hematocrit (Hct) in 2014 and 2015. All plotted relationships (model-based regression lines; individual points based on the model output) are significant ($P<0.001$), except FL vs. PL in 2015 ($P = 0.933$).

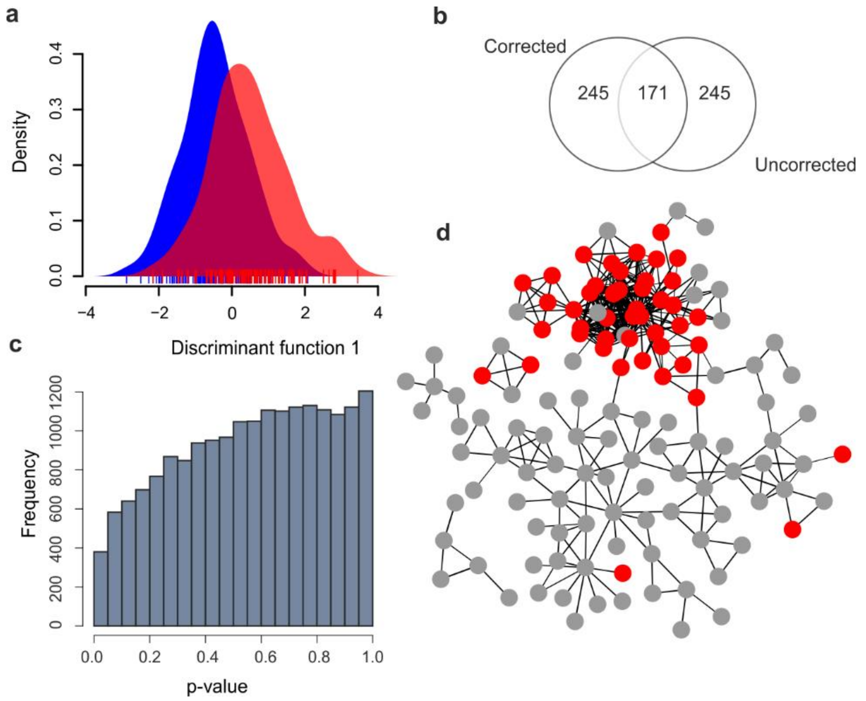


**Fig. S3. Differential gene expression analyses using uncorrected survival estimates.** (**a**) Density distribution of the first discriminant scores correspond to non-survivors and survivors (blue and red areas, respectively); (**b**) p-value distribution from differential gene expression analysis is hill-shaped, rather than uniform, when using uncorrected survival estimates, indicating that misclassification of individuals likely resulted in a violation of statistical test assumptions; (**c**) Venn diagram showing overlap among 416 top genes identified from differential expression analysis for corrected and uncorrected survival; (**d**) protein–protein interaction network based on 416 top genes correlated with uncorrected survival. Mitotic cell cycle genes (GO:0000278, FDR = $8.6 \times 10^{-17}$, n = 57) in the protein–protein interaction network are shown as red circles.

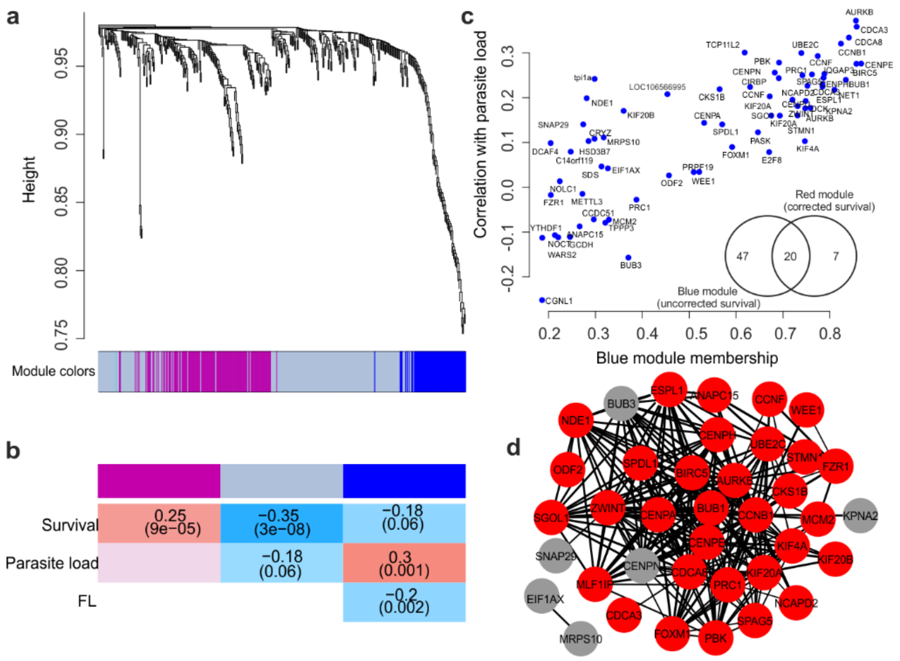


**Fig. S4. Weighted gene co-expression network analysis of top 416 genes correlated with uncorrected survival and their relationship with parasite load and fork length.** (**a**) Gene dendrogram with the corresponding two modules. Each color represents a module with highly connected genes; (**b**) Relationships of module eigengenes and survival, parasite load and fork length (FL). Numbers in the table show correlation coefficients between the corresponding module eigengene and trait, with the p-values in brackets; (**c**) Module membership of the genes in the blue module and the corresponding correlation coefficients with parasite load. The inset Venn diagram illustrates the overlap between the blue module (uncorrected survival) and the red module (corrected survival; Fig. 3); (**d**) Protein-protein network of the genes in the blue module linked to mitotic cell-cycle (GO:0000278; n = 35; FDR = $1.85 \times 10^{-30}$) are shown as red circles.

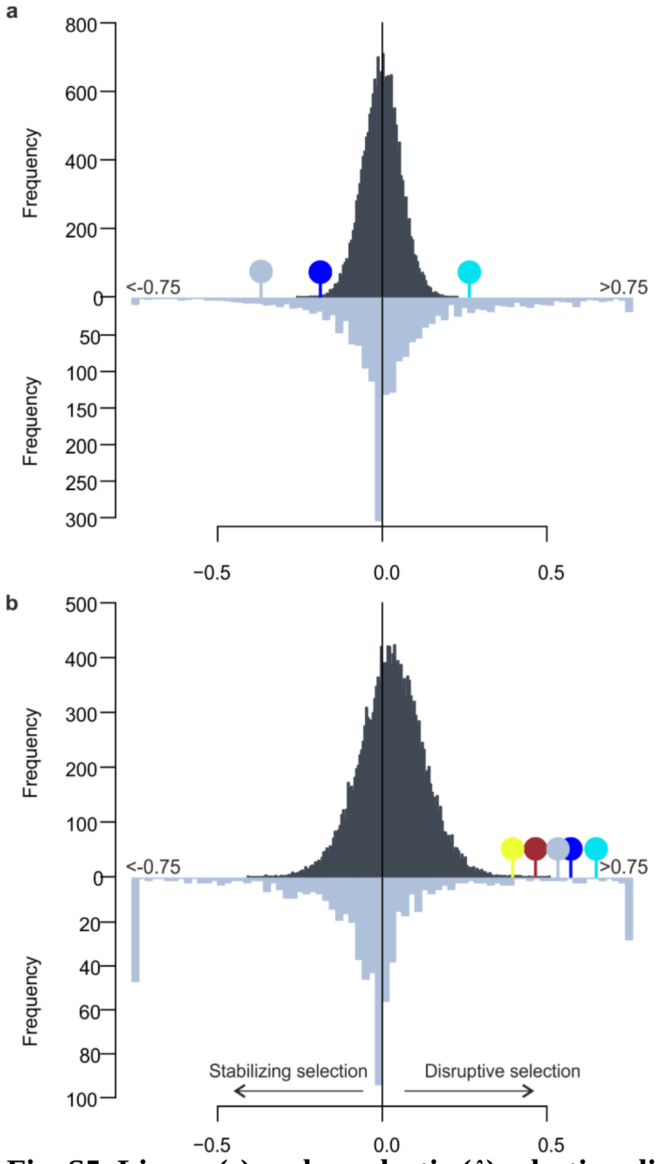


**Fig. S5. Linear (*s*) and quadratic (*λ*) selection differential estimates** based on uncorrected survival for 18,717 transcripts (dark grey histogram, above the line) and published phenotypic traits (*4*) (light grey histogram, below the line). Selection differentials for the WGCNA gene modules are shown as colored pins. (**a**) Linear selection differentials s; (**b**) Quadratic selection differentials λ. The distribution of quadratic selection coefficients λ was strongly shifted towards the right tail (two-sample Wilcoxon test, $P = 6.2 \times 10^{-09}$).

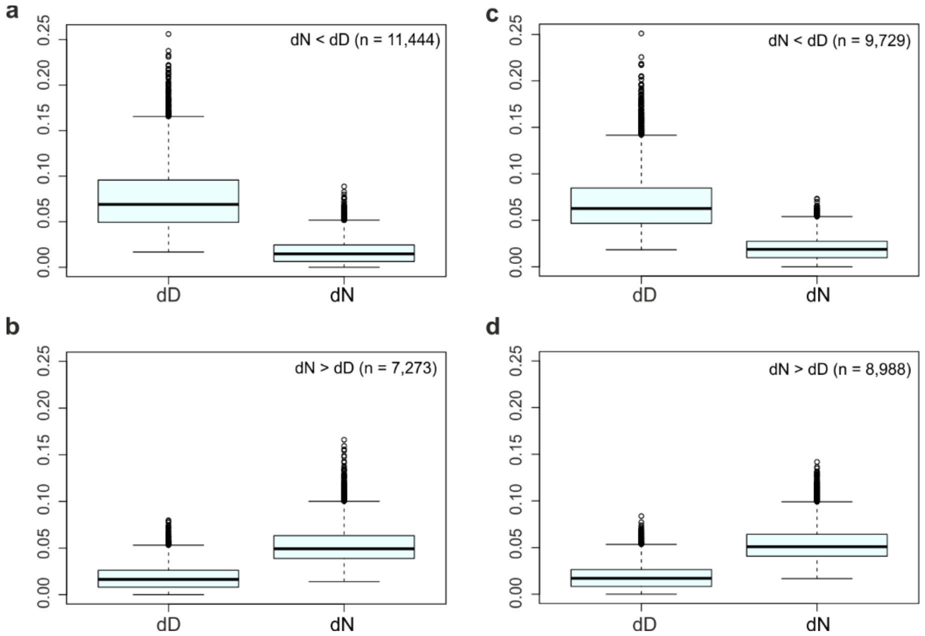


**Fig. S6. Components of the distributional selection differential** (*23*) representing selection on the trait mean (dD = |s|) and on the shape of the trait distribution (dN), calculated based on corrected (**A**, & **B**) and uncorrected (**C** & **D**) survival estimates. The line across the box represents the median, the box edges represent the upper and lower quartile, the whiskers extend to a maximum of 1.5 × IQR beyond the box, and the points represent outliers.

**Table S1. Genetic mark-recapture results.** Numbers in bold indicate captured individuals in September (Area 9 corresponds to the uppermost location), non-bold numbers in Catch II and III column reflect inferred estimates. The five main target areas (1-5) were electrofished thrice, whereas the remaining areas were fished once in September. Estimated numbers of survived but uncaptured fish are based on mean capture probability derived from depletion electrofishing (*35*) and the area-specific frequency of recaptured fish.

| Area ID | Initial sampling date | Initial catch | Recapture sample date | Catch I | Catch II | Catch III | No. of recaptures | Recapture frequency | Capture probability (95%CI) | Census size (95%CI) | Estimated no. of uncaptured fish | Estimated no. of uncaptured survivors that were sampled in August |
|---|---|---|---|---|---|---|---|---|---|---|---|---|
| -1 | | | 25.9.2015 | **26** | 7.7 | 3.8 | 0 | 0.00 | 0.65 (0.60-0.70) | 39.2 (38.1-40.3) | 13.1 | 0.0 |
| 0 | | | 25.9.2015 | **30** | 8.9 | 4.4 | 8 | 0.27 | 0.65 (0.60-0.70) | 45.2 (44.0-46.5) | 15.2 | 4.0 |
| 1 | 30.8.2015 | 46 | 22.9.2015 | **39** | **5** | **6** | 10 | 0.20 | 0.71 (0.58-0.85) | 51.0 (48.2-53.8) | 1.0 | 0.2 |
| 2 | 30.8.2015 | 48 | 22.9.2015 | **60** | **18** | **6** | 26 | 0.31 | 0.70 (0.59-0.81) | 86.0 (82.0-90.0) | 2.0 | 0.6 |
| 3 | 30.8.2015 | 72 | 24.9.2015 | **61** | **27** | **11** | 27 | 0.27 | 0.58 (0.45-0.70) | 107.0 (97.4-116.6) | 8.0 | 2.2 |
| 4 | 30.8.2015 | 52 | 24.9.2015 | **62** | **16** | **10** | 30 | 0.34 | 0.65 (0.53-0.76) | 92.0 (86.2-97.8) | 4.0 | 1.4 |
| 5 | 30.8.2015 | 60 | 25.9.2015 | **85** | **25** | **12** | 42 | 0.34 | 0.66 (0.56-0.75) | 127.0 (120.6-133.4) | 5.0 | 1.7 |
| 6 | | | 25.9.2015 | **48** | 14.2 | 7.0 | 3 | 0.06 | 0.65 (0.60-0.70) | 72.4 (70.4-74.4) | 24.3 | 1.5 |
| 7 | | | 25.9.2015 | **53** | 15.7 | 7.8 | 2 | 0.04 | 0.65 (0.60-0.70) | 79.8 (77.7-82.2) | 26.8 | 1.0 |
| 8 | | | 27.9.2015 | **64** | 19.0 | 9.4 | 1 | 0.02 | 0.65 (0.60-0.70) | 96.3 (93.8-99.2) | 32.3 | 0.5 |
| 9 | | | 27.9.2015 | **22** | 6.5 | 3.2 | 1 | 0.05 | 0.65 (0.60-0.70) | 33.1 (32.3-34.1) | 11.1 | 0.5 |

**Table S4. Number of survivors and nonsurvivors.** Corrected 3'-RNAseq dataset is based on RF analysis, which identified a small number of individuals (n = 13) that possessed transcriptomic signatures characteristic to survivors but were not re-captured in September.

| Category | Survivors | Nonsurvivors |
|---|---|---|
| Initial mark-recapture (n = 278) | 150 (54%) | 128 (46%) |
| Uncorrected 3'-RNAseq dataset (n = 238) | 114 (48%) | 124 (52%) |
| Corrected 3'-RNAseq dataset (n = 238) | 127 (53%) | 111 (47%) |
